# Linking single-cell transcriptomes with secretion using secretion-encoded single-cell sequencing (SEC-seq)

**DOI:** 10.1101/2024.05.17.594711

**Authors:** Justin Langerman, Sevana Baghdasarian, Rene Yu-Hong Cheng, Richard G. James, Kathrin Plath, Dino Di Carlo

## Abstract

Cells secrete numerous proteins and other biomolecules into their surroundings to achieve critical functions – from communicating with other cells to blocking the activity of pathogens. Secretion of cytokines, growth factors, extracellular vesicles, and even recombinant biologic drugs defines the therapeutic potency of many cell therapies. However, gene expression states that drive specific secretory phenotypes are largely unknown. We provide a protocol that enables linking the Secretion amount of a target protein EnCoded (SEC) by thousands of single cells with transcriptional sequencing (seq). SEC-seq leverages microscale hydrogel particles called Nanovials to isolate cells and capture their secretions in close proximity, oligonucleotide-labeled antibodies to tag secretions on Nanovials, and flow cytometry and single-cell RNA-sequencing platforms for readout. Cells on Nanovials can be sorted based on viability, secretion amount, or other surface markers without fixation or permeabilization, and cell and secretion-containing Nanovials are directly introduced into microfluidic droplets-in-oil emulsions for single-cell barcoding of cell transcriptomes and secretions. We have used SEC-seq to link T-cell receptor sequences to the relative amount of associated cytokine secretions, surface marker gene expression with a highly secreting and potential regenerative population of mesenchymal stromal cells, and the transcriptome with high immunoglobulin secretion from plasma cells. Nanovial modification and cell loading takes under 4 hours, and once the desired incubation time is over, staining, cell sorting, and emulsion generation for scRNA-seq can also be completed in under 4 hours. By linking gene expression and secretory strength, SEC-seq can expand our understanding of cell secretion, how it is regulated, and how it can be engineered to make better therapies.

## Introduction

Cells secrete numerous biomolecules that facilitate many organismal processes such as inflammatory stimulation, neuronal signaling, chemotactic recruitment of other cell types to a location, extracellular matrix (ECM) scaffolding of tissue, developmental signaling to establish spatial differentiation programs, and maintenance of niche supporting adjacent cells. Examples of crucial secretory functions span developmental biology, immunology, and regeneration; including regulation of developmental gradient morphogens for patterning of nearly all organs in early embryos via secretion of bone morphogenic protein and sonic hedgehog, targeting and neutralization of pathogens via secreted antibody from plasma B cells, and priming for wound healing and regeneration by collagen and vascular endothelial growth factor secretion from fibroblasts and mesenchymal stromal cells (MSCs).

Secreted proteins make up approximately 25% of the proteome in mammalian cells^1^. The rate and capacity of protein secretion can depend on numerous trafficking and storage steps within the cell before release (e.g. granzyme B, perforin and associated molecules are stored in cytotoxic granules which are primed for release upon cytotoxic T cell activation). Gene transcription may therefore be a poor proxy for ultimate protein secretion levels, and biochemical methods for directly measuring secretion from individual cells have lagged behind those for nucleic acid sequencing^2^. Single-cell transcriptomics have revealed that an astounding heterogeneity exists even within cell samples considered to be one cell type. Protein secretions are also released in a heterogeneous manner, so consideration of secretions at a single-cell level is required to capture the full scope of secretion biology and the expected functional heterogeneity of secretions from a cell population/product. Understanding the links between RNA transcription and protein secretions at the single-cell level can better elucidate the biological processes that control secretion and can reveal new insights by overlaying functional secretion information on populations of cells that are transcriptionally defined. Secreted proteins are also often used in biotechnology; for production of recombinant products, biologic drugs, and as secreted therapeutics or growth factors from cells for various downstream applications. Therefore, a better understanding of the transcriptional underpinnings for robust, long-term secretion can drive increases in yields and stability of biotechnology products.

### Development of the protocol

To link the amount of secretion of a target protein from a cell with the same cell’s transcriptome we have developed Secretion EnCoded single-cell sequencing, or SEC-seq^3–5^. As a first step, we needed to devise a method by which the secretions from a single cell could be spatially associated with it in a downstream transcriptomic workflow. To achieve spatial localization of secretions, we used microscale hydrogel particles (Nanovials)^6,7^. Nanovials are cavity-containing hydrogel particles that have localized surface chemistry for functionalization with affinity reagents such as extracellular matrix molecules or antibodies that bind cells and also can capture the secreted protein of interest^8^ (Figure 1). Individual cells can be loaded into Nanovials by using a ratio of cells to Nanovials that leads to 10-20% of Nanovials being cell-loaded. In this situation, cells are loaded into Nanovials following Poisson statistics, with the majority of cell-loaded Nanovials containing a single cell^6^. After loading of cells into the Nanovials, the cells remain viable and can be incubated for short or long time periods to capture and accumulate secretions, with up to 24 hours tested. We next developed a method by which the secreted protein amount could be linked with the transcriptomic data for each individual cell. As in the Cellular Indexing of Transcripts and Epitopes (CITE-seq)^9^ protocol for linking cell-surface proteins, we used a secondary oligo-barcode labeled antibody against the secretion product of interest, forming a sandwich (Figure 1). This effectively translates the quantity of secretion to an oligonucleotide signal that can be quantified by standard single-cell RNA sequencing (scRNA-seq) workflows. To make the protocol accessible, we designed the Nanovials to be compatible with commercial scRNA-seq devices, specifically, the 10x Genomics Chromium instrument and microfluidic droplet generator chips. The cell-loaded Nanovials are small enough to be loaded directly into these devices, so Nanovials can be first sorted by fluorescence-activated cell sorting (FACS) to remove unloaded Nanovials and then be directly introduced into the single-cell sequencing platform (Figure 1). Using standard single cell sequencing workflows, libraries for quantification of linked RNA and secretions are created, completing the SEC-seq workflow. This enables the analysis of a secretion of interest in the full context of the single-cell transcriptomic information for 1000s of individual cells.

**Figure 1.**
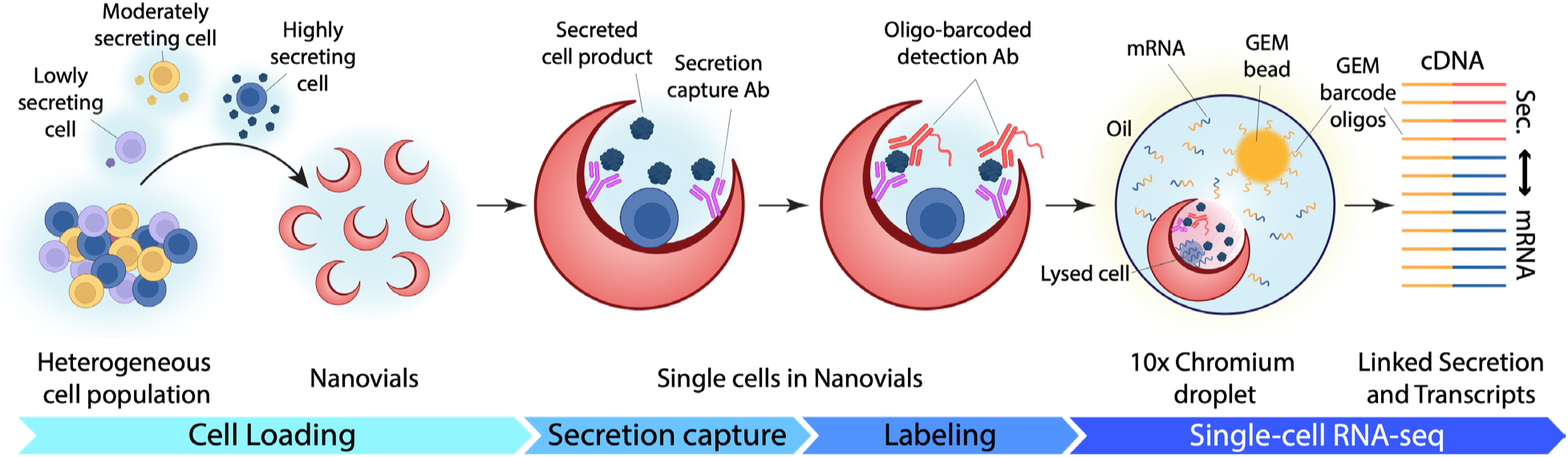
– A summary of the Secretion EnCoded single-cell sequencing (SEC-seq) workflow. Cells are loaded into Nanovials and their secretions are captured by attached antibodies. The secretion levels are quantified utilizing an oligo-barcoded secretion detection antibody at a single-cell level and are linked to the transcriptomes of each cell through emulsification of the cell-loaded Nanovials into droplets using standard single-cell sequencing workflows. Schematic partially created with bioRENDER (www.biorender.com).

### Applications of the method

To date, SEC-seq has been used in three different applications where secretions and gene expression data were important to link (Figure 2). In plasma cells, SEC-seq was used to investigate which cells were high secretors of immunoglobulin G (IgG), an antibody secretion required for fighting off infections and pathogens^3^ (Figure 2a). IgG secretion was linked to single cell transcriptomes for 10,000 cells. SEC-seq confirmed that several known plasma cell markers were in fact associated with high IgG secretion, and found new surrogate markers for highly secreting plasma cells. This study defined several transcriptional features of highly secreting plasma cells, including increased expression of oxidative phosphorylation genes, MYC target genes, and interferon response genes. SEC-seq is enabling the development of potential therapeutics to modulate IgG secretion in diseases, to either increase or decrease as might be helpful.

**Figure 2.**
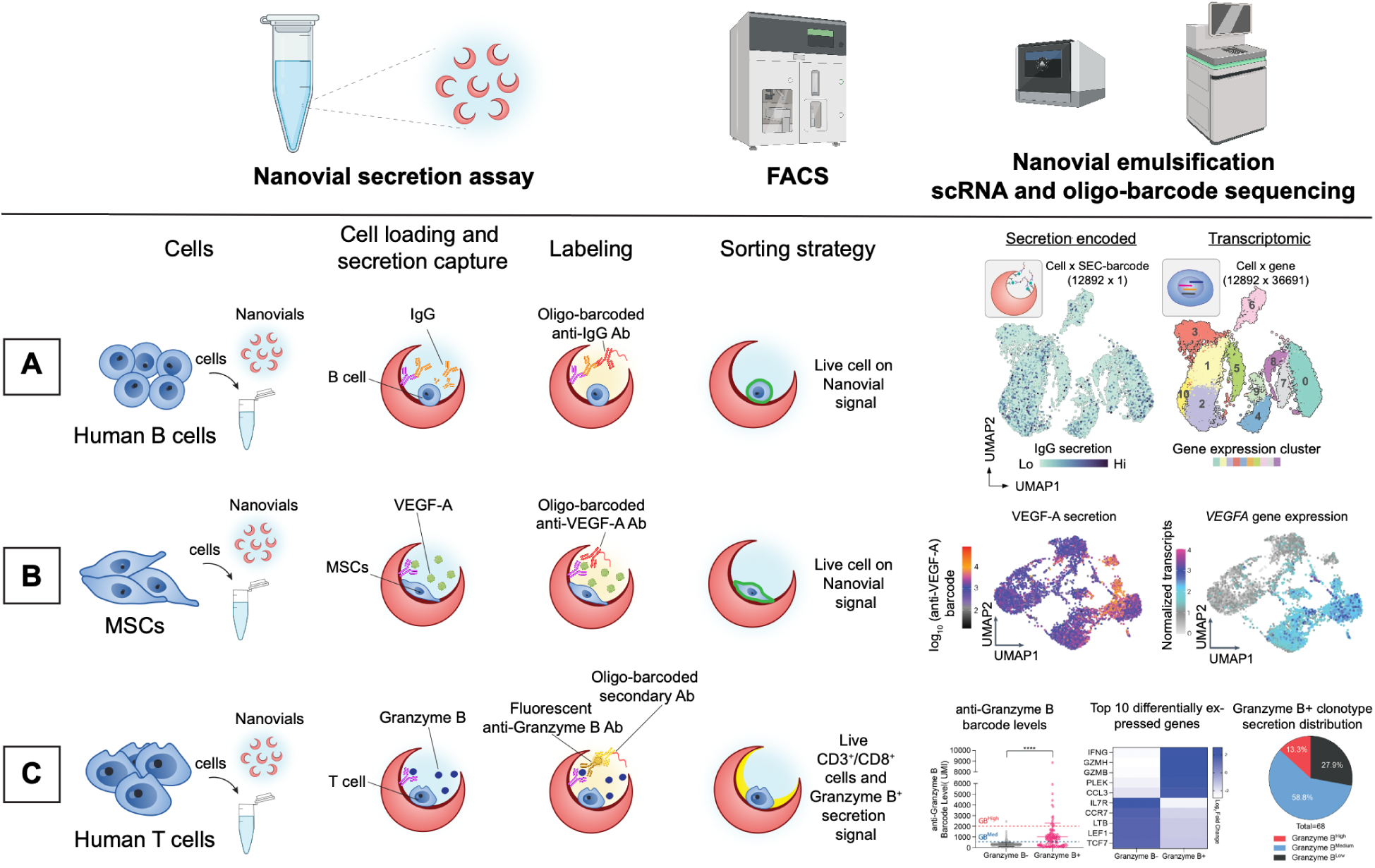
Demonstrations of SEC-seq with three distinct cell types and secretions and the resulting data. The amount of A) IgG secreted by Human B cells and B) VEGF-A secreted by Mesenchymal stromal cells were measured and linked to the transcriptomes from the same cells. C) The amount of granzyme B secreted by Human T cells were measured and linked to V(D)J sequences where TCR sequences were ranked based on the corresponding cell’s secretion level. Schematic partially created with bioRENDER (www.biorender.com).

Another well-known secretion, the endothelial promoting factor, Vascular Endothelial Growth Factor A (VEGF-A), was also analyzed by SEC-seq in mesenchymal stromal cells (MSCs), to determine the connection between transcriptional identity and VEGF-A secretion, a measure for the functional regenerative potential of MSCs^4^ (Figure 2b). By linking the transcriptome to VEGF-A secretion it was discovered that VEGF-A secretion is heterogeneous and that transcription of the *VEGFA* gene has very low correlation with the secretion level of the protein. Intriguingly, a particular subpopulation of the MSCs dominates VEGF-A secretion, which was not predictable by *VEGFA* transcript levels. It was possible to define a specific gene expression profile of this subpopulation, which included higher baseline transcription of many secretory factors and specific surface proteins. VEGF-A therefore stands out as an example for an important secretion product that cannot be described solely by its transcriptional levels. Instead, SEC-seq revealed that a phenotypically secretory and pro-regenerative subpopulation of MSCs secrets VEGF-A highly without modifying regulation of the gene itself.

Another powerful example of SEC-seq was its use in characterizing T cells isolated by their T-cell receptor (TCR) specificity. T cells expressing TCRs that recognize specific antigenic peptides presented by major histocompatibility complex I (pMHC) could be isolated by linking the pMHC molecules on Nanovials and then characterizing the correlated secretion of effector molecules triggered for secretion from the specifically bound T cells^5^ (Figure 2c). Captured cells were then interrogated by SEC-seq for granzyme B secretion, which was linked to both their transcriptomes and their specific TCR. The sequences of these unique TCRs could also be recovered allowing for future genetic manipulation. It was also found that TCRs from cells with higher granzyme B secretion were more likely to be functional when transduced into new cells than TCRs associated with lower granzyme B levels. SEC-seq allowed for the isolation of TCRs and increased confidence in the correct matching between pMHC and TCR sequence, a task which has proven to be a fundamentally challenging previously due to the diversity of TCRs expressed.

Beyond these published applications, the SEC-seq workflow is modular, enabling a researcher to investigate any secretion provided affinity reagents are available. The technique can also be multiplexed for the detection of several secreted products^5^, or even measure other types of communication factors released by cells, such as extracellular vesicles (EVs)^10^. The approach is equally suited for adherent and suspension cells through binding to the extracellular matrix and/or antibodies specific to cell surface markers. SEC-seq can probe biomolecules secreted at high rates (like IgG) or low rates (like VEGF-A) by adjusting the time allotted for secretion accumulation. Secretions induced by cell-cell interactions can also be analyzed, as demonstrated with T cells secreting cytokines in response to interaction with pMHC molecules bound to the Nanovial surface^5^. Finally, we anticipate other techniques, like functional approaches such as Perturb-seq can be combined with SEC-seq to delve into the network of genes controlling secretion of various factors.

### Comparison with other methods

There are only a few methods available to measure the secretions of single cells^11^, and only in the last year have techniques been introduced that link secretion and transcriptomes of single cells at scale. Traditionally, intracellular levels of secreted proteins, like cytokines, were measured in immunology studies, as a proxy for cellular secretion. In these techniques cells are fixed and permeabilized and then stained with antibodies against the target protein. It is understood that intracellular stores of proteins may not reflect what is secreted, and the process of fixation and permeabilization also is known to degrade mRNA, making downstream transcriptomic measurements noisy^12^. Other techniques, such as ELISpot and the Isolight system (now part of Bruker) enable measurement of proteins secreted from the cell, which are captured on surfaces surrounding cells and detected with fluorophore-or enzyme-labeled antibodies^13^. However, these techniques do not link secretion with transcriptomic information. In linking with transcriptomes, SEC-seq most directly resembles CITE-seq, an oligo-barcoded antibody method that quantifies surface protein abundance alongside RNA^9^. SEC-seq is instead adapted for secretion capture within the local solid-phase surfaces provided by modified Nanovials. Previous attempts used microwell chips to capture secretions in proximity to single cells performing sandwich immunoassays, and hand picking cells one by one for single-cell RNA-sequencing^14^. Because of the laborious nature of the protocol only a few dozens of cells could be analyzed. Concurrently with the development of SEC-seq, another secretion and transcriptome method called Time Resolved Assessment of single cell Protein secretion (TRAP-seq)^15^ was published that similarly uses an oligo-barcoded antibody to detect secretions in a typical scRNAseq workflow. This method uses a custom hybrid bi-specific antibody, anchoring the primary secretion capture antibody to the cell using covalent linkage to a second antibody targeting a highly-expressed surface protein on the cell of interest. This then requires a third detection antibody against the secretion. TRAP-seq is dependent on development of the custom bi-specific antibody, uniformly– and highly-expressed surface antigens, and methods to prevent cross-talk from cells secreting in proximity to each other as secretions are not contained within a cavity. SEC-seq relies on Nanovials which capture secretions over the larger area of the Nanovial cavity and prevent convective transport which can reduce cross-talk^16^. The larger Nanovial surface also provides more flexibility in secretion incubation time, secretion multiplexing, and cell type compatibility with both adherent and suspension cells. The solid phase of the Nanovial also enables presentation of molecules which can trigger secretions in a cell-type specific manner, as demonstrated by the binding and activation of T cells to pMHC-coated Nanovials.

### Expertise needed to implement the protocol

One of the key advantages of the protocol is that all instruments and reagents used are commercially available. Biotinylated and matrix coated EZM™ Nanovials are available in 35 and 50 µm varieties from Partillion Bioscience. The handling of Nanovials, attachment of antibodies, and loading of cells into Nanovials requires no special equipment or expertise beyond needed for routine handling of the cell type of interest. Most steps are accomplished via pipetting, centrifugation, and incubation. Cell sorting and scRNA-seq workflows require access to FACS and scRNA-seq instruments and basic expertise in flow cytometry and single-cell RNA-sequencing; both aspects can be handled by core facilities if available. Some expertise in analyzing scRNA-seq data sets is suggested or bioinformatics core facilities can assist with data analysis.

### Limitations

Nanovial loading and dynamic range of secretions are limited by the size of the Nanovial. Loading into Nanovials requires dissociation into single cells and a cell size that can fit into a Nanovial cavity. Nanovial size is limited by compatibility with the scRNA-seq workflow, which for the 10x Genomics Chromium instrument and microfluidic chips was 35 micrometers on the outer diameter in our previous experiments^3–5^. However, microfluidic chips have recently become available from 10x Genomics (GEM-X chips) that are expected to be compatible with larger Nanovials (50 micrometers or greater). Additionally, other single-cell RNA-sequencing technologies that rely on split-pool reactions and do not use microfluidic chips are also expected to be compatible^17–19^. Smaller Nanovials have less binding sites for secreted products, likely leading to reduced dynamic range compared to the larger Nanovials and limiting the multiplexing of secretion targets due to competition for the binding sites for secretion products.

Cell loading onto Nanovials should be optimized. Loading may require troubleshooting or the use of a cell-capture antibody, depending on cell type. It should also be considered that the Nanovial environment can alter the baseline expression state of cells. We observed that MSCs loaded on Nanovials had altered gene transcription, as might be expected for any perturbation in growth conditions^4^. Viability, subpopulations captured, and differentiation state, or ability to differentiate were not affected^4^. The upregulated genes were related to cell division, which is not expected to affect the majority of biological questions.

Current Nanovial formulations are biotinylated, which requires streptavidin-biotin chemistry to perform any functionalization. This limits the assays for cell and/or secretion product capture to antibodies or other affinity agents that are biotinylated. Since only a single moiety (biotin) is available for functionalization, if multiple functionalization steps need to be performed, such as different antibodies for cell and secretion capture, then these should all be performed at the same time. Because many types of media contain biotin, we recommend functionalization of Nanovials prior to loading cells which are in biotin-containing media. This may result in the limitation that pre-functionalized Nanovials are prone to non-specific secretion capture from unloaded cells during the cell loading procedure.

It is important to remember that SEC-seq measures the secretion product that accumulates in Nanovials over the entire incubation period. Secretion may not occur uniformly over a long time period and these temporal differences are not detected without SEC-seq time courses. Because of these and other temporal dynamics, there may be a fundamental mismatch between the cumulative secreted protein and the transcriptomic snapshot taken at the end of the experiment, which should be considered when interpreting results. Depending on the level of the secretion, cross talk is possible between loaded Nanovials, as unbound secretions can diffuse out of the Nanovial. We have shown that this background signal is orders of magnitude lower in empty Nanovials, but true zero values will be obscured^4^. Finally, detection of secretions is limited to available sandwich antibody pairs with the correct modifications (biotin and oligo-attachment).

### Experimental design

In principle, SEC-seq should be broadly applicable to any cell type that can be loaded into the Nanovial cavity. Setup should begin with optimizing cell loading. Once conditions are determined, Nanovials are modified and loaded with cells, then allowed to secrete during an incubation time. Prior to emulsion generation, cell-loaded Nanovials are typically enriched using FACS (through the use of a cell surface marker, viability stain, secretion amount, or a combination thereof). Nanovials can be loaded directly into a variety of cell sorters^20^(www.partillion.com/protocols). The wide availability of these devices indicates you can apply the protocol exactly as follows.

Following enrichment of loaded Nanovials, the protocol described here uses a 10x Genomics scRNA-seq machine and reagents, a commonly available commercial solution to create single-cell cDNA and oligo-barcode libraries. The secondary antibody used for detection of secretions requires modification with an oligonucleotide barcode. The protocol described here leverages the 10x Genomics Feature Barcode primer site on the oligo primer gel beads used in single-cell sequencing, which can capture oligo barcodes such as the TotalSeq™ barcodes (Biolegend). SEC-seq has been tested on the 10x Genomics single-cell workflows. Deviation may require redesign, especially in the case of using a different scRNA-seq method. In principle SEC-seq will be compatible with any scRNA-seq method, but the oligo-barcoded antibody detection design will have to be reconsidered. Using the recommended setup, SEC-seq should be easily adapted to any cell type and any secretion.

### Preparation of the Nanovials and antibodies

EZM™ Nanovials are commercially available from Partillion Bioscience with functionalized biotin sites within the interior cavity. This allows for the addition of new modifications that can be attached via the streptavidin-biotin interaction. In a normal SEC-seq workflow, this usually includes attachment of an antibody that can recognize the secretion prior to cell loading, via a short incubation. Secretion target capture is based on antibody recognition, using a primary and secondary antibody similar to enzyme-linked immunosorbent assays (ELISAs) or ELISpot assays. Any secreted protein which has an ELISA pair available is already primed for use in SEC-seq. The primary capture antibody must be ordered with a biotin moiety, which is commonly available from various vendors. The detection antibody should be labeled with a barcode amenable with the chosen scRNA-seq workflow. It is also possible to use an epitope-tagged detection antibody, such an anti-target-APC or PE-tagged antibody. In this case an oligo-tagged secondary antibody against that epitope (e.g. barcoded anti-APC antibodies) can be used. If using the 10x Genomics platform with support for Feature Barcode capture sites, one can modify the detection antibody with TotalSeq™ B barcodes by ordering reagents from Biolegend. The conjugation of an antibody with the oligo-barcode can take upwards of two months in development time if you are using a primary tagged detection antibody. Using a secondary oligo-barcoded detection antibody against a primary detection antibody is also a viable strategy to obtain readily available reagents for your target secretion. For example, in the T-cell study^5^, granzyme B was targeted with by an APC-conjugated antibody, and detection was mediated by a secondary oligo-barcoded anti-APC antibody.

### Loading the Nanovials

Adherent cells can often be loaded directly onto the primary gelatin substrate within the cavity of EZM™ Nanovials, due to native expression of integrins which bind the Nanovial’s gelatin substrate. Non-adherent cell types may require modification of the Nanovial with a cell capture antibody that is specific to a surface protein constitutively expressed in the cell type of interest, which has been shown to capture non-adherent cells in the Nanovial cavity at high levels^3,5,6,21^. Alternatively, other affinity agents, like antigens, or pMHC molecules can be loaded on Nanovials to load antigen-specific cells. Cells should be tested for their binding prior to a secretion capture experiment. When loading, using more Nanovials increases the probability of a given cell being loaded, increasing the fraction of cells bound, but leading to more empty Nanovials. Loading should start at 1 cell: 1 Nanovial to decrease the chance of multi-cell loading. Loading generally follows Poisson statistics, with 1:1 loading yielding 10-20% of Nanovials with single cells. For rarer material, the amount of Nanovials can be increased to 1 cell: 3-5 Nanovials to ensure higher cell recovery (although lower percentage of Nanovials with cells). We suggest loading at a ratio that leads to at least ∼10% of nanovials containing a cell, which you can determine via simple microscopy or flow cytometry. This fraction will be enriched later before emulsion generation in the single-cell library preparation, since empty Nanovials are removed during a cell sorting step. Loading should be checked by applying a fluorescent cell stain such as Calcein AM and checking via flow cytometry or by counting manually on a microscope.

### Testing the dynamic range of secretion capture

It is advised to test several facets of the secreted protein capture and detection on Nanovials prior to a SEC-seq experiment using an analogous flow cytometry-based assay, especially if you are developing SEC-seq against a new secreted analyte. Use a fluorescent antibody against the target secretion for assay calibration. First, use a dilution series of recombinant protein of the secretion of interest, incubated with empty Nanovials containing the secretion capture antibody. Then bind the fluorescent detection antibody and use flow cytometry to confirm a dose dependent response in fluorescent signal. Next, a trial using the same fluorescent detection schema should be run on live loaded cells to determine that the secretion signal is detectable. Previous studies have found secretion signal heterogeneity from secreting cells that ranges over two orders of magnitude^4^. Gating on loaded Nanovials via cell staining can confirm that the secretion is localized to cell-loaded Nanovials and the crosstalk to empty Nanovials (and therefore overall) is minimal at this step. A fluorescence detection assay can also be used to determine the best incubation time for the SEC-seq experiment, as secretions can vary in release intensity from saturation at 1 hour for some proteins to continual accrual at 24 hours for others^3,4^. Cells are capable of surviving and dividing in the Nanovials, with new cells eventually budding out of a full cavity. This may be a consideration for long incubation times.

### Secretion and transcriptome capture

After loading and incubating cells for an optimal secretion time, the oligo-barcoded detection antibody is added. Oligo-barcoded detection antibody can directly bind to the target secretion or to a fluorophore (APC, PE) bound to a primary detection antibody bound to the target secretion. It is strongly advised to enrich for cell-loaded Nanovials and discard multi-cell loaded Nanovials before proceeding to scRNA-seq, using a cell stain and FACS. To load cell-containing Nanovials into the 10x Genomics Chromium workflow, each Nanovial is treated as a cell and Nanovials are counted and diluted to the ‘cell’ concentration of the relevant single-cell sequencing approach. Do not overload Nanovials in the 10x Genomics workflow; although this practice is common for certain scRNA-seq strategies, overloading of Nanovials can cause clogging. We recommend targeting 10,000 “cells” or fewer for recovery.

Nanovials can be loaded directly into the 10x Genomics Chromium flow chips in a similar manner as large cells. The faster sedimentation of Nanovials compared to cells can lead to uneven distribution of Nanovials into the emulsion. To avoid this, it is recommended to use a modified 10x Genomics chip loading strategy as detailed in the protocol. This should allow the Nanovials to enter the emulsion with a more regular distribution and help prevent clogging or sedimentation out of the flow. SEC-seq has been tested on 10x Genomics Chip G flow devices, although we also expect improved compatibility with the larger GEM-X chips.

### Analysis

Transcriptomic data obtained by SEC-seq can be analyzed via normal methods with no special consideration^22^. The yield of transcripts of cells on Nanovials is roughly comparable to free cells^4^. The secretion detection antibody information is contained in the Feature Barcode data if the 10x Genomics workflow is used and can be extracted as a per cell value of secretion. We have found this metric does not correlate with transcriptome depth, and therefore do not find it appropriate to normalize the secretion signal using that approach. Thousands of reads of the secretion detection antibody barcode are detected per cell, typically, so a log transformation is used for working with the secretion data.

Possible approaches for analysis include assigning per cluster or per cell type secretion levels to determine which cells are the primary secretors, comparison to known secretor or non-secretor data as control cells, correlation of secretion to transcript levels and possible gene networks to determine potential causal relationships, and measurement of the heterogeneity of secretion within cell populations delineated by the scRNA-seq data.

## Reagents and Equipment

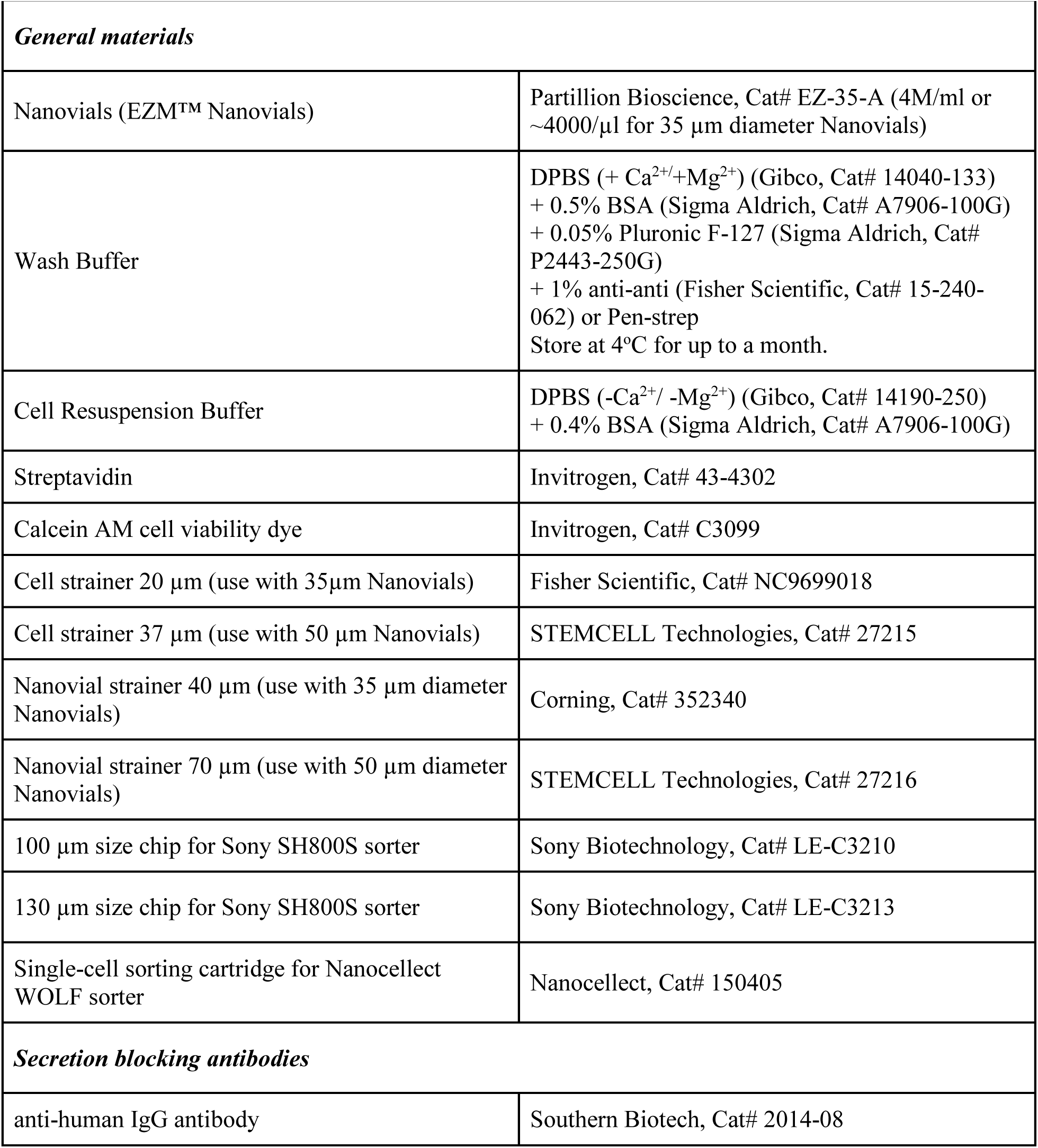
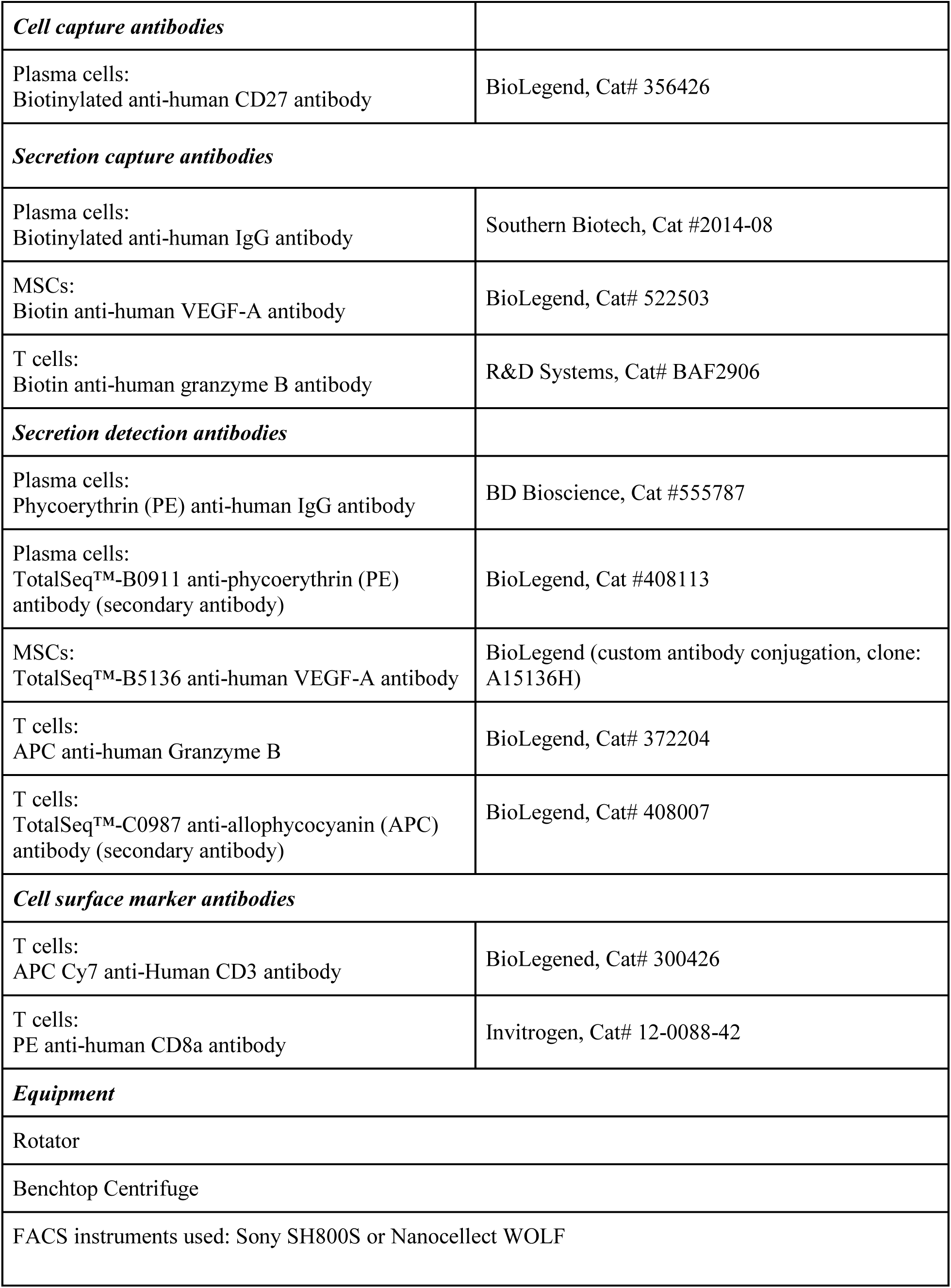
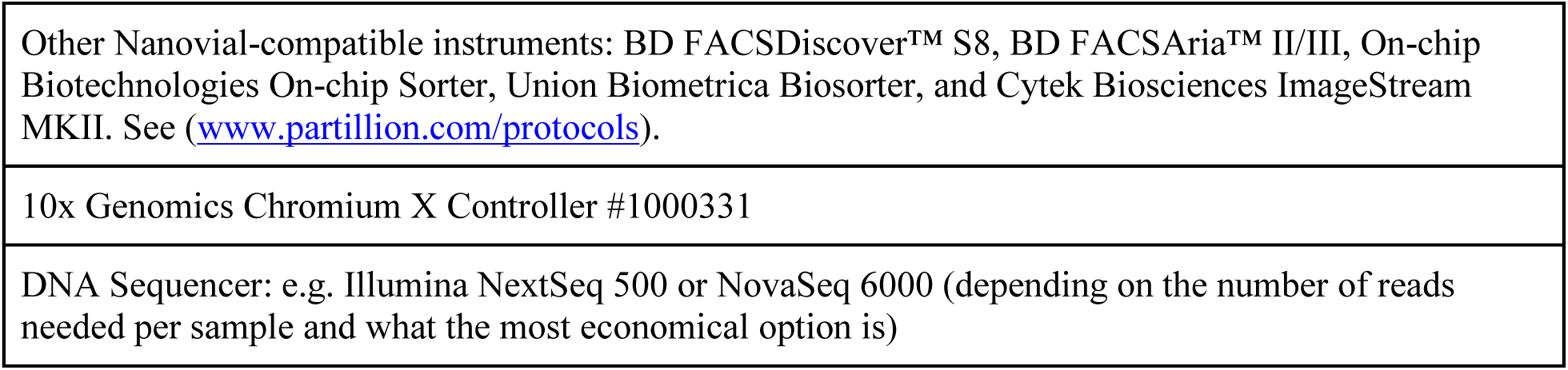

## Procedure

These steps are also outlined in Figure 3.

**Figure 3.**
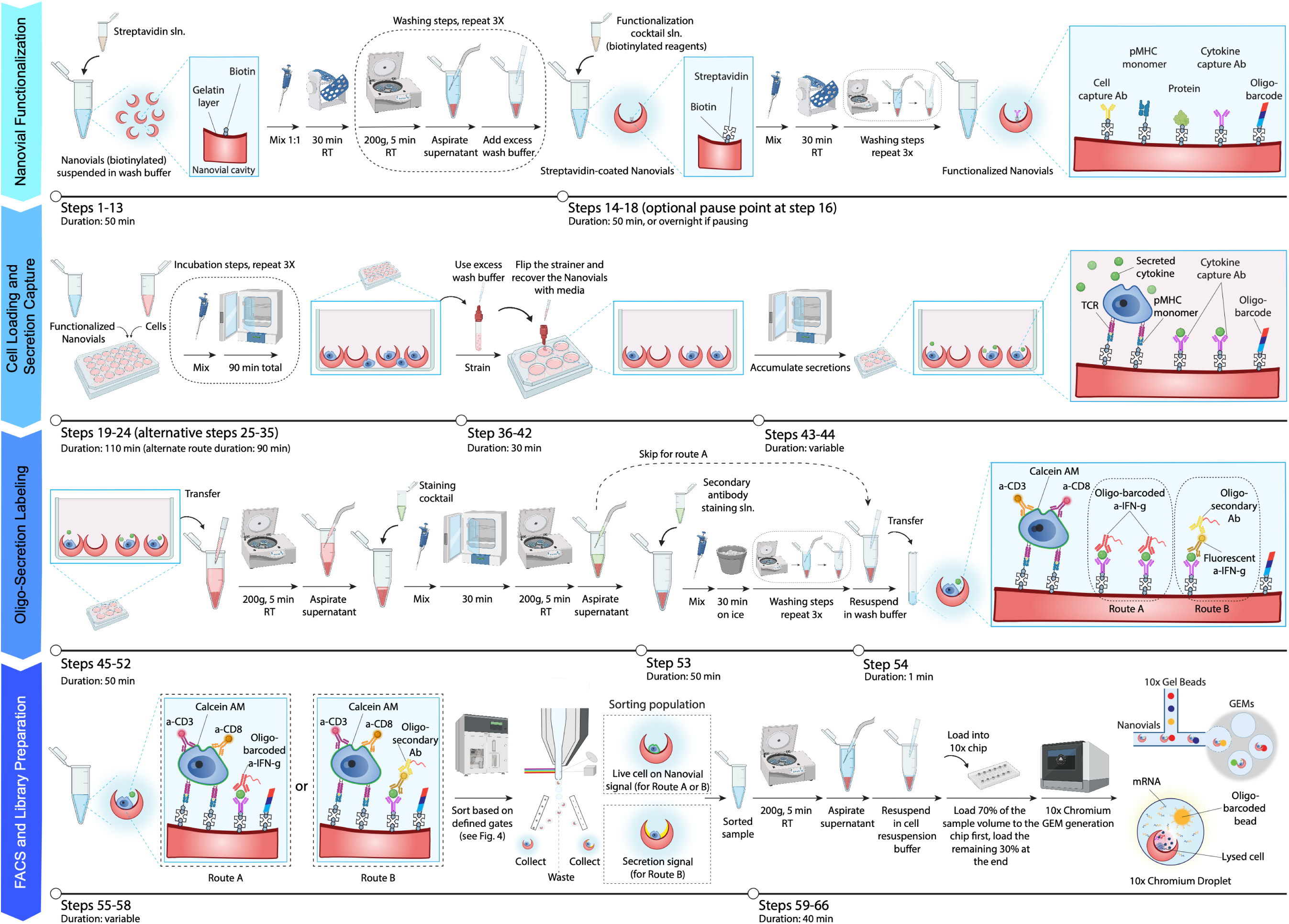
– Detailed SEC-seq protocol. Nanovials are first functionalized with cell and secretion capture moieties. Cells are suspended in media and mixed with Nanovials in a tube or well plate followed by straining to remove unbound cells. Cell-loaded Nanovials are incubated to collect secretions which are then stained with oligo-labeled detection antibodies. The sample is then sorted utilizing either live cell and/or secretion signals for sorting, followed by single-cell sequencing. Schematic partially created with bioRENDER (www.biorender.com).

Nanovial coating (Volumes are based on 200k Nanovials per vessel) Timing: Steps 1–13, 50 min; Steps 14-18, 50 min, or overnight if pausing;

1. Confirm the Nanovial and cell numbers to use for loading. For one SEC-seq experiment, with a target of 10k-40k cell-loaded Nanovials use ∼200k Nanovials. Multiple vessels with this 200k increment can be processed in parallel to scale the experiment, to test multiple conditions or larger cell numbers. The stock concentration of Nanovials is ∼4000 Nanovials/µl for 35 µm diameter Nanovials and ∼2000 Nanovials/µl for 50 µm diameter Nanovials. The cell-to-Nanovial ratio should be 1:1 to load ∼10-20% of Nanovials with cells, accounting for loss in the loading process.

## Note on Cell Loading

The number of Nanovials used will depend on the loading rate for the cells of interest. Lower cell-to-Nanovial ratios (e.g. 1:5 or 1:10) can be used for rare cell samples to isolate more of the inputted cells into Nanovials, and to reduce doublets for smaller cells. However, this will result in a smaller fraction of Nanovials loaded with cells.

2. Precoat all pipette tips used to handle Nanovials with Wash Buffer by pipetting the maximum volume up and then expelling the volume.
3. Precoat a 1.5 ml eppendorf tube with 500 µl of Wash Buffer by flushing the surface and removing the volume.

## Note

Precoating reduces the loss of Nanovials that may stick to the pipette tip and vessel surfaces.

4. Pipette the Nanovial stock to resuspend equally throughout the volume and make sure there is no visible pellet.
5. Pipette the Nanovial suspension from the stock and transfer it to a precoated 1.5 ml eppendorf tube with a precoated tip. The volume should be determined based on the number of Nanovials (ultimately cells) that are desired to be analyzed (see Step 1). For example, pipette 50 µl from the 4000/µl stock to transfer 200k 35-µm-diameter Nanovials sufficient for 10k-40k loaded cells.
6. In a separate tube, prepare an equal volume of streptavidin solution at a concentration of 300 µg/ml in Wash Buffer.
7. Transfer the streptavidin solution to the tube containing the Nanovial suspension.
8. Mix thoroughly by pipetting up and down (for at least 30 seconds) and incubate for 30 min, RT.
9. Add a 10X volume of Wash Buffer to the reaction.
10. Centrifuge at 200g for 5 mins, RT.
11. Aspirate supernatant being careful not to aspirate the Nanovial pellet.
12. Repeat wash steps 9-11, three times to completely wash off any unbound streptavidin.
13. Resuspend Nanovials back into 1 volume (ex. 50 µl) of Wash Buffer.
14. Prepare a Capture Cocktail; this cocktail can include (i) a biotinylated surface marker-specific cell capture antibody (or other biotinylated cell binding agents) at a final concentration of 50 µg/ml and (ii) a biotinylated secretion capture antibody at a final concentration of 50 µg/ml in 1 volume of Wash Buffer (ex. 50 µl).

## Note

Creating the Capture Cocktail is a necessary step since both cell capture and secretion capture moieties bind to the same streptavidin functional groups on the Nanovials. Sequential addition will lead to low binding of the subsequently added capture moiety.

15. Add 1 volume of Capture Cocktail to the streptavidin-coated Nanovial suspension from step 13.
16. Incubate for 30 min at RT or 4°C overnight if pausing the experiment. PAUSE POINT: The Nanovials mixed with Capture Cocktail can be kept overnight at 4°C and the experiment can be resumed at the next step the following day. This is recommended for secretion durations of longer than 1 hour to reduce the overall work duration in a single day.
17. Add 10X volume of Wash Buffer to the Nanovial suspension from step 16.
18. Spin at 200g for 5 min and aspirate the supernatant, being careful not to aspirate the Nanovial pellet. Repeat the wash steps three times total.

### Cell loading

Timing: Steps 19-24, 110 min;

19. Optional: For cells with low viability (below 70%), it is recommended to remove dead cells by live cell enrichment or Ficoll gradient separation prior to loading into Nanovials.
20. Resuspend cells in a 20X volume of media to a final concentration such that there is 1 cell per 1 Nanovial (ex. 200k cells in 1 ml per 200k Nanovials in 50 µl).
21. Precoat either a 24-well plate or a 5 ml eppendorf tube with 2 ml of Wash Buffer per well or tube.
22. Add the entire volume (ex. 50 µl) of Nanovial suspension into a single well or tube from step 21.
23. Add cell suspension into the well or tube and pipette up and down gently for 30 seconds in circular motions to mix cells and Nanovials.
24. Incubate the plate or tube at 37°C for 90 min, with agitation (e.g. 20 rpm) if possible. Pipette up and down gently to mix every 30 min.

## Note

Secretions, such as immunoglobulins, that occur rapidly at high levels may lead to high background during the standard 90 min cell loading process. In this case, use this Cell loading process for high secretion rate cells instead, which uses blocking and cooling to limit background.

### Cell loading: High-secretion-rate cells (Alternative to Steps 19-24)

Timing: Steps 25-35, 90 min;

25. Place the Nanovial suspension from step 18 on the ice and place 20 ml cell culture medium on ice to equilibrate temperature.
26. Transfer target number of cells in culture medium into a 15 ml falcon tube.
27. Centrifuge cells at 4°C at a speed suitable for the cell type.
28. Wash cells with 15 ml ice cold culture medium and centrifuge again at 4°C and a speed suitable for the cell type.
29. Resuspend cells in cell culture medium at a concentration of 4000k cells/ml (50 µl for 200k cells).
30. Optional: For high secretion rate cells, it may be beneficial to add blocking antibodies against the secreted protein in order to reduce or eliminate the capture of secreted proteins during cell loading steps. Therefore, dilute the blocking antibody (stock concentration 0.5 mg/ml) in culture medium at a final concentration of 50 µg/ml into the cell suspension. The blocking antibodies against the secreted protein should be the same clone as the secreted protein capture antibody.
31. Pipette to mix the cell suspension (50 µl) with blocking antibody (7.7 µl) 3 times.
32. Transfer the cell suspension (57.7 µl) into the Nanovial suspension tube (∼20 µl) from step 18 on ice. Blocking antibody is ∼10-fold diluted in step 31-32.
33. Pipette up and down gently for 30 seconds in circular motions to mix cells and Nanovials evenly.
34. Add 1 ml ice cold cell culture medium per 200k Nanovials.
35. Incubate tube on ice for 1 hr allowing cells to adhere to Nanovials.

### Removing unloaded cells

Timing: Steps 36-42, 30 min;

36. Set aside either two 15 ml conical tubes, or one 15 ml conical tube and one 6-well plate. One 15 ml conical tube is for collection of waste (unloaded cells). The second 15 ml conical tube or the 6-well plate is for collecting Nanovials.

## Note

A 6-well plate should be used for longer secretion incubation times (> 90 minutes) to maintain normal culture conditions during incubation, or when a larger fraction of cells are expected to secrete, as the larger area of the plate spaces cell-loaded Nanovials further apart.

37. Precoat one of the 15 ml conical tubes or a 6-well plate with Wash Buffer and set aside (for collecting loaded Nanovials).
38. Precoat a 20µm cell strainer with Wash Buffer by pipetting 500 µl of Wash Buffer on top of the strainer and letting it flow through.
39. Place the cell strainer on top of another 15 ml conical tube (for unloaded cells), with the narrow end facing up (inverted compared to normal usage).
40. Transfer the Nanovials and cell mix onto the strainer and let the liquid drain through the strainer.
41. After the sample flows through, wash at least two more times by pipetting 500 µl of Wash Buffer on top of the strainer and letting it flow through.
42. Flip the strainer over and recover the Nanovials into either the precoated 15 ml conical tube or 6-well plate by pipetting 2 ml of media and letting it drain through the inverted strainer.

### Incubation to collect secretions on Nanovials

Timing: Steps 43-44, timing depends on the cell secretion rate (e.g., 30 min for plasma cells secreting IgG, 3-6 hours for T cells secreting cytokines, 12-24 hours for stromal cells secreting VEGF-A).

43. For normal secretion studies, incubate at 37°C in the well plate or tube. Time varies by secretion, 1-24 hours.
44. Optional: For high-secretion-rate cells, spin down Nanovials at 200g for 5 min after straining, resuspend strained Nanovials with 2 ml pre-warmed culture medium per 200k Nanovials and transfer into a pre-coated 5 ml eppendorf tube. Load the tube on a rotator/shaker at low speed at 37°C and incubate for 30 min to collect secretions.

### Labeling secretions with detection antibodies

Timing: Steps 45-52, 50 min; Step 53, 50 min; Step 54, 1 min;

45. Precoat a 5 ml eppendorf tube with Wash Buffer and leave 2 ml of Wash Buffer in the tube.
46. Transfer the Nanovial suspension from step 43 or 44 to the tube.
47. Centrifuge samples at 200g for 5 min, RT, and aspirate the supernatant to wash.
48. Prepare detection Staining Cocktail in Wash Buffer for labeling secreted proteins and cell surface markers. Typical dilutions are 1:250 to 1:2000 for 0.5 mg/mL antibody stocks in 100 µl of Wash Buffer. For IgG oligo-barcoded detection antibodies (TotalSeq™ B), this was 0.5 µl in 1 ml for a 1:2000 dilution, for the VEGF-A oligo-barcoded detection antibody this was 1:700 to a final concentration of 71.5 µg/ml.

## Note

The Staining Cocktail should also contain a cell stain to enable selecting cell-loaded Nanovials by FACS. Option 1: Add viability dye to the Staining Cocktail. For calcein AM, add 0.2 µl of stock per 1 ml. Option 2: For heterogeneous cell populations, add fluorescent antibodies against cell surface markers of target cells. For anti-CD3, 1:30 of 100 µg/ml stock.

49. Add 100 µl of the Staining Cocktail solution to the 5 ml tube containing Nanovials from step 47, pipette to mix gently, and incubate at 37°C for 30 min. Optional: For high-secretion-rate cells, stain on ice for 20 min.
50. Repeat steps 51-52 two times to wash samples with 1 ml of Wash Buffer.
51. Centrifuge samples at 200g for 5 min, 4°C.
52. Aspirate supernatant, being careful not to aspirate the Nanovial pellet. If no secondary staining is desired, proceed to step 54 (route A).
53. Optional (route B) – If using a secondary secretion detection antibody for either oligo-barcoding or fluorescence-based sorting, repeat staining steps as from step 49-52 for secondary reagents. This secondary staining can also occur after FACS for addition of an oligo-barcoded secondary antibody targeting the primary fluorescent secretion detection antibody.
54. Resuspend the pellet obtained from either step 52 or 53 in 300 µl Wash Buffer and transfer to a FACS tube for sorting.

### Sorting

Timing: Steps 55-58, variable, depending on the samples and sorting rate of the machine;

55. Optional – For 35 µm Nanovials, filter samples with a 40 µm strainer to remove clumps. For 55 µm Nanovials, filter samples with a 70 µm strainer instead.
56. Choose appropriate flow cell or microfluidic chip size compatible with Nanovials (100 µm nozzle for 35 µm Nanovials or 130 µm nozzle for 35 or 50 µm Nanovials). See de Rutte et al.^20^. The smaller nozzle size of 100 µm is preferred for the 35 µm Nanovials as it enables sorting at higher rates.
57. Set up the gates for live single-cell-loaded Nanovials, and/or distinguishing cells of interest (based on surface markers or secreted markers) loaded onto the Nanovials (see Figure 4).
58. Collect sorted Nanovial samples for the scRNA-seq workflow into a 2ml eppendorf tube pre-filled with 500 µl of Wash Buffer.

**Figure 4.**
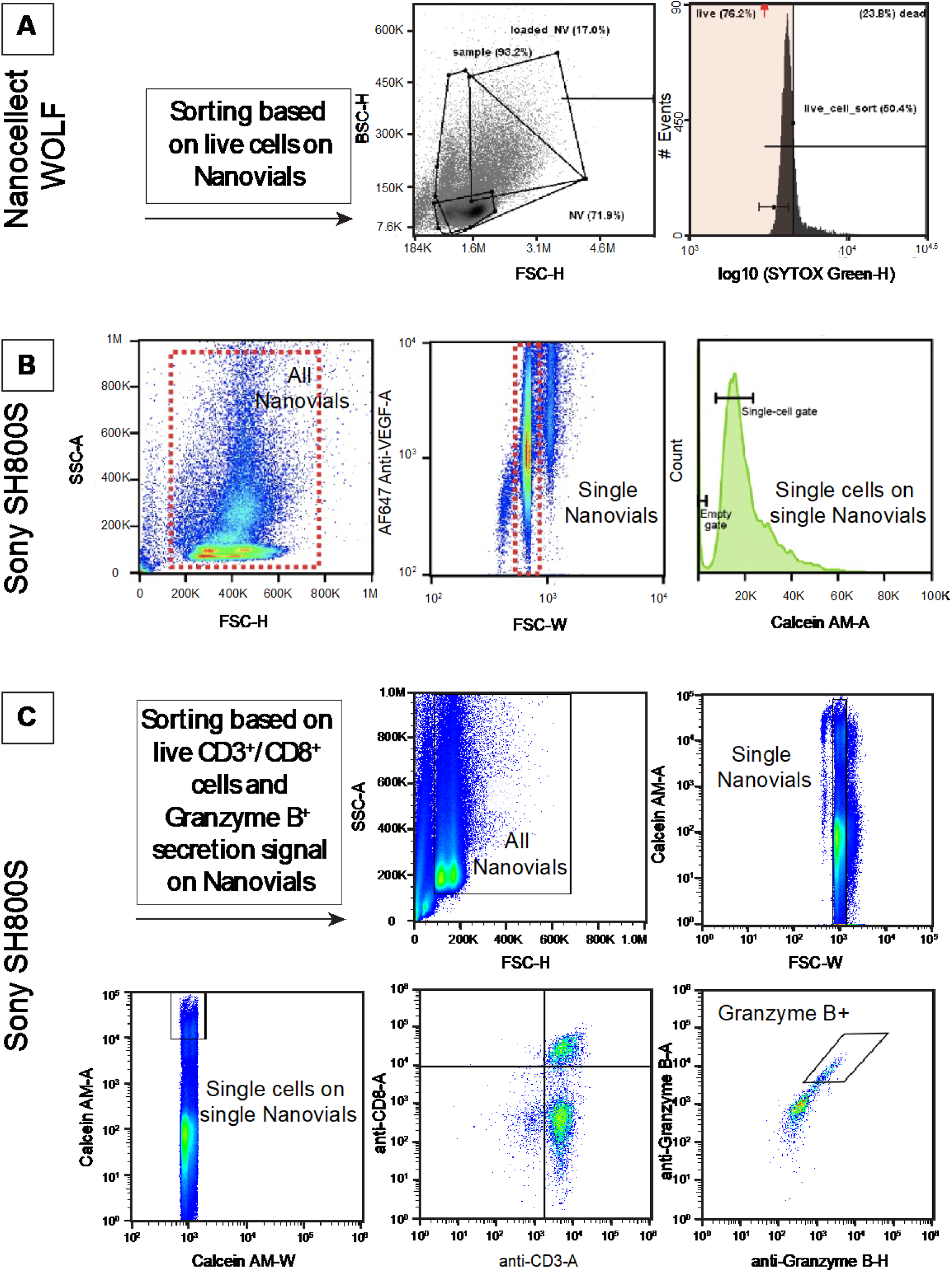
– Gating strategies for sorting single-cell loaded Nanovials for sequencing and analysis. A) Plasma cells sorted with the Nanocellect WOLF sorter based on Nanovial scatter and live cell staining. B) Human MSCs sorted with the Sony SH800S sorter based on single Nanovial scatter and single live cell staining. C) Human T cells sorted with the Sony SH800S sorter based on single Nanovial scatter, live CD3+/CD8+ cells, and high Granzyme B secretion signal on Nanovials.

### 10x Genomics scRNA-seq workflow, library preparation and sequencing

Timing: Steps 59-66, 40 min; Steps 67-68, 4-7 days;

59. Calculate Nanovial number targeting loading of 2.5k-10k Nanovials per sample for each 10x Genomics chip well.
60. Centrifuge Nanovials at 200g for 5 min at 4°C.
61. Resuspend Nanovials in Cell Resuspension Buffer such that the final concentration is 500 Nanovials/µl.
62. Prepare 10x Genomics workflow as usual. Make a mixture of the Nanovial suspension and Reverse Transcriptase solution. Prepare this mixture just before loading of the Chip G/X. Please refer to the “Cell Suspension Volume Calculator” in the 10x Genomics user handbook to determine the optimum final volume for the Nanovial suspension based on concentration.
63. Allow Nanovials to sediment in Reverse Transcriptase mixture during reagent preparation (∼3 min). Take approximately 70% of sample volume supernatant without disturbing Nanovials and load into the cell well. This is ∼50 µl in the Chip G workflow.
64. Load all other wells with appropriate reagents.
65. Return to the remaining 30% of Nanovial/Reverse Transcriptase mixture, resuspend and load over the top of the cell well volume and attach the gasket (approx. 20 µl in Chip G workflow)
66. Proceed immediately to the controller and start flow.
67. Prepare libraries using 10x Genomics Chromium Next GEM Single Cell v3.1 kit following the 10x Genomics User Guide (CG000317) as usual. Be sure to also prepare the Feature Barcode library.
68. Sequencing the libraries using the following minimum read lengths for a 138 cycle sequencing run: 28 bp Read1, 10 bp i7 Index, 10 bp i5 Index, and 90 bp Read2. Refer to 10x Genomics User Guide for library sequencing recommendations.

### SEC-seq data analysis

69. Demultiplex any indexed data and convert to FastQ.
70. Process Fastq files using the Cellranger multi function to process both the transcriptome and secretion oligo-barcoded detection antibody reads. Providing the appropriate barcodes in the multi configuration file allows for correct assignment of detected capture antibody oligo barcodes to each cell at this step.
71. Perform data analysis including normalization, batch correction, dimensional reduction, hierarchical clustering, clustering, and differential gene analysis using typical workflows. Secretion values can be extracted from the raw data. Python: See https://github.com/Rene2718/SEC-seq_plasma-cell_nanovial R: Using Seurat, read the Cellranger output with the Read10x() function, secretion data per cell will be output in a slot of the created object, named as in your Cellranger configuration settings.

## Troubleshooting

See Figure 5 for a troubleshooting workflow.

| Step | Problem | Possible reason | Solution |
| --- | --- | --- | --- |
| 18 | Aggregation of Nanovials | Too high centrifuge speed during washing steps.<br>Incomplete or uneven mixing of streptavidin or other antibody cocktail solutions with Nanovials. | Spin Nanovials only at 200g. Coat all vessels and tips with Wash Buffer solution prior to manipulation to prevent aggregation. Pipette or flick tube to disperse Nanovials, rather than vortexing. Carefully and gently mix reagents enough times. |
| 24 | Poor cell loading / mostly empty Nanovials | Low expression of target marker targeted by cell capture antibody. Cell capture antibody is not binding to the cells or is at too low concentration on Nanovial. Insufficient cell loading time. Excess pipetting or handling leading to cell loss. | Try a different cell capture antibody clone or target. Increase time for cell loading. Switch from 37°C loading to loading on ice or vice versa. Apply intermittent agitation (every 15 to 30 min) during loading incubation, rotation or gentle shaking. |
| 54 | No secretion signal | Insufficient time to accumulate secretions. Secretion capture antibody does not binding secretion or is too low concentration on Nanovial. | Troubleshoot reagents using recombinant protein sweeps and fluorescent antibodies. Confirm cell secretion does happen with ELISPOT. Try longer time points to allow for more secretion. |
| 54 | High secretion signal on all empty Nanovials | High-secreting cells secrete during the loading step, many unbound cells secreting leading to high background.<br>Too long incubation time during the secretion step.<br>Detection or capture antibody cross species reactivity, e.g., capture anti-IgG (human) binds detection antibodies (mouse IgG). | Use blocking antibodies during cell loading steps and load on ice to reduce cell secretion. Tune secretion time for cell type, and analyte. Disperse Nanovials in a well of a well plate instead of tube. Replace antibodies with those having minimum reactivity to other species or with pre-adsorbed antibodies. |
| 66 | No emulsion forms during single-cell library construction | Obstruction of single-cell sequencing chip. Time between Nanovial loading and running chip was too long. | Run the chip immediately after loading. Sort for single Nanovials. Use smaller Nanovials. Use a larger, GEM-X chip. |
| 67 | No antibody oligo library during preparation | May have used wrong primers at first cDNA amplification step during library prep. | Make sure to use Feature barcode workflow/primers that capture antibody conjugated oligo-barcode during cDNA amplification. |
| 71 | Low quality scRNAseq data | Cells may have died or were lost from Nanovials prior to scRNAseq workflow. | Confirm viability and Nanovial loading by fluorescence microscopy prior to and after sorting with FACS. |

**Figure 5.**
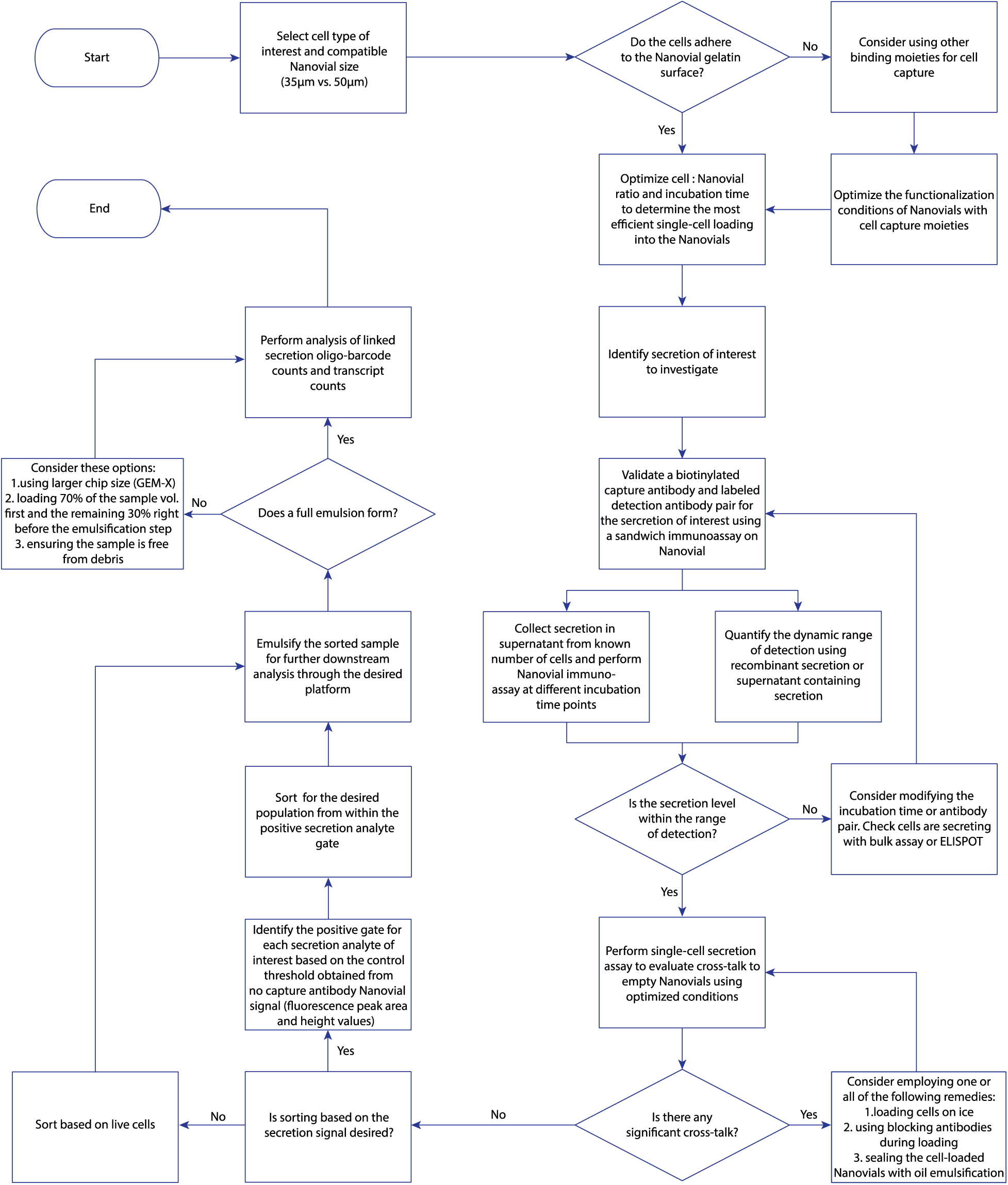
– Workflow for geting started and troubleshooting for SEC-seq. A first-time user can follow the flowchart to plan, validate, and troubleshoot a SEC-seq experiment for a desired application.

## Anticipated results

Loading of cells is cell-type dependent, but when starting with a ratio of 200k cells:200k Nanovials we expect 20k-40k cells to be loaded, sufficient for obtaining up to 10k final cell-loaded nanovials for scRNA-seq. If a cell type does not bind well to the matrix on the Nanovials, loading of less cells may be observed which suggests that an antibody affinity agent should be used. Using that method, in theory, any type of cell can be loaded into the Nanovials. Previous experiments have used 35 µm or 50 µm diameter Nanovials, which should be compatible with cells having diameters of <20 µm or 30 µm respectively. Cells can be loaded and survive in the Nanovials for days and can even divide and “bud” out of the Nanovials if enough time passes. Sorting based on viability dye quantity can limit analysis to single cells if this is a concern. Cells have normal viability after recovery from Nanovials. Gene expression is slightly altered by loading into Nanovials, but cell type differences persist and the affected genes seem to be in pathways related to maintaining attachment and cell division when compared to cells in suspension.

Development of an assay for a secretion of interest using antibody reagents should be relatively straightforward. For any particular secreted product, an expected distribution of fluorescence signal targeting the secretion above a no-capture control suggests you may proceed to SEC-Seq. Some secretions we targeted failed to have a differential distribution versus the control; in these cases, either the antibody pair is not working, or the secretion product is not meaningfully secreted. If differential secretion is picked up via flow cytometry, then that is usually reflected by similar meaningful differences in the SEC-seq data.

FACS enrichment for single-cell-containing Nanovials and collection and counting should have result in recovery of 50-80% of sorted events. Nanovials are easy to spin and recover in the various buffer changes before scRNA-seq. To confirm correct emulsion generation in scRNA-seq, immediately after initiating flow a small amount of emulsion can be optionally sampled from the outlet well to visualize Nanovials within the droplets. In the event of low emulsion volume, there may have been a clog on the 10x Genomics chip (see troubleshooting section). SEC-seq should yield data for a number of cells just slightly below typical 10x Genomics scRNA-seq experiments, roughly in the 80-90% range. Libraries are constructed and sequenced using normal methods. The cDNA library from Nanovials will have the same size ranges as free cells, the feature barcode library for the capture barcode is typically ∼220 nt. We generally targeted 10k cells with 1k Nanovials/µl, which yielded 3-4k single-cell transcriptomes and matched secretion feature barcode reads. Reads can be processed normally using any single-cell analysis method, the Cellranger workflow perfectly suits this approach. For the secretions tested in the noted publications, thousands of capture antibody barcodes per cell were usually detected. The heterogeneity and specificity were dependent on the particular secretion. We found no correlation between a cell’s transcript depth and secretion amount in our studies, so for normalization only a log transformation is recommended to flatten the distribution for display.

## Author contributions statements

J.L. S.B. and R.C. developed the methodology and wrote the manuscript. J.L. and S.B. contributed writing to all sections, constructed the figures, and edited the manuscript. K.P. and R.J. contributed guidance for the writing and edited the manuscript. D.D. contributed writing to all sections, edited the manuscript, and helped design figures.

## Acknowledgments

This project has been made possible in part by grant 2023-332386 from the Chan Zuckerberg Initiative Donor Advised Fund (CZI DAF), an advised fund of the Silicon Valley Community Foundation.

Selected schematic figures were partially created using bioRENDER (www.biorender.com).

